# Genetic loci with parent of origin effects cause hybrid seed lethality between *Mimulus* species

**DOI:** 10.1101/022863

**Authors:** Austin G. Garner, Amanda M. Kenney, Lila Fishman, Andrea L. Sweigart

## Abstract

The classic finding in both flowering plants and mammals that hybrid lethality often depends on parent of origin effects suggests that divergence in the underlying loci might be an important source of hybrid incompatibilities between species. In flowering plants, there is now good evidence from diverse taxa that seed lethality arising from interploidy crosses is often caused by endosperm defects associated with deregulated imprinted genes. A similar seed lethality phenotype occurs in many crosses between closely related diploid species, but the genetic basis of this form of early-acting F_1_ postzygotic reproductive isolation is largely unknown. Here, we show that F_1_ hybrid seed lethality is an exceptionally strong isolating barrier between two closely related *Mimulus* species, *M. guttatus* and *M. tilingii*, with reciprocal crosses producing less than 1% viable seeds. Using a powerful crossing design and high-resolution genetic mapping, we identify both maternally- and paternally-derived loci that contribute to hybrid seed incompatibility. Strikingly, these two sets of loci are largely non-overlapping, providing strong evidence that genes with parent of origin effects are the primary driver of F_1_ hybrid seed lethality between *M. guttatus* and *M. tilingii*. We find a highly polygenic basis for both parental components of hybrid seed lethality suggesting that multiple incompatibility loci have accumulated to cause strong postzygotic isolation between these closely related species. Our genetic mapping experiment also reveals hybrid transmission ratio distortion and chromosomal differentiation, two additional correlates of functional and genomic divergence between species.

## INTRODUCTION

Understanding the genetics of interspecific incompatibility provides insight both into the origins of species barriers and the processes driving genomic divergence within species. In plants, postmating incompatibilities are often significant, and in particular, many early plant geneticists noted difficulties in generating F_1_ hybrids from experimental crosses between closely related species (Thompson 1930, Stebbins 1957, Valentine and Woodell 1963, Vickery 1978). Some of these crossing problems are likely caused by pollen-pistil interactions that prevent fertilization altogether, but for others, successful fertilization occurs only to end in a high rate of F_1_ hybrid seed lethality. This failure of F_1_ seeds is also a common outcome of interploidy crosses (known as triploid block, Ramsey and Schemske 1998), and represents a form of postzygotic reproductive isolation that acts early in the life cycle. Relative to other forms of hybrid incompatibility (*e.g*. hybrid necrosis or hybrid sterility), which may act in older plants or only in one sex, F_1_ hybrid seed lethality might be a particularly strong barrier to interspecific gene flow. Moreover, its prevalence in plant crosses from diverse taxa suggests that F_1_ hybrid seed failure might represent a major class of reproductive isolation in plants – albeit one that has gone largely unmentioned in modern accounts of speciation (*e.g*., Coyne and Orr 2004, but see Tiffin and Moyle 2001, Turelli and Moyle 2007).

Hybrid seed inviability has long been thought to result from developmental defects in the endosperm, an important nutritive tissue for the developing embryo (Brink and Cooper 1947, Woodell and Valentine 1961). In the seeds of flowering plants, a triploid endosperm is one of the two products of double fertilization. The male gametophyte (pollen tube) releases two sperm into the ovule: one fuses with the egg nucleus to produce the zygote, and the other fuses with the two nuclei of the central cell to form the endosperm. Classic work showed that deviations from the ratio of two maternal to one paternal genomes (2m:1p) disrupts endosperm development (Johnston *et al.* 1980), explaining the failure of many interploidy crosses, which double the maternal or paternal contribution. A similar disturbance to endosperm “balance” also seems to cause hybrid seed lethality in many diploid crosses between species (Valentine and Woodell 1963, Johnston and Hanneman 1982, Josefsson *et al.* 2006, Ishikawa *et al.* 2011).

The sensitivity of the endosperm to mismatches in parental genome dosage led to the hypothesis that genomic imprinting is the molecular cause of both interploidy and interspecific seed lethality (Haig and Westoby 1991, Birchler 1993, Gutierrez-Marcos 2003). The expression of an imprinted gene is dependent on its parent of origin due to differential epigenetic modifications established during male and female gametogenesis (Kohler *et al.* 2012). If imprinted genes encode dosage-sensitive regulators, quantitative changes in parental contributions could cause a stoichiometric imbalance with downstream targets (Birchler *et al.* 2001, Birchler and Veitia 2012). This mechanism provides an explanation for the common observation that reciprocal interploidy crosses differ in seed phenotypes (*e.g*., Thompson 1930, Brink and Cooper 1947, Scott *et al*. 1998, Sekine *et al*. 2013); disruptions to maternally versus paternally derived alleles have different consequences for endosperm development. Recently, several studies in *Arabidopsis* have shown that imprinted genes are misexpressed in interploidy crosses (Erilova *et al.* 2009, Jullien and Berger 2010, Wolff *et al.* 2011, Kradolfer *et al*. 2013). In principle, similar types of deregulation involving imprinted genes might occur in the endosperm of F_1_ hybrid seeds between diploid species (Kohler *et al.* 2010), but aside from two studies in *Arabidopsis* (Josefsson *et al.* 2006, Burkart-Waco *et al*. 2012) and *Capsella* (Rebernig *et al*. 2015), this idea remains largely untested.

Intriguingly, parent of origin effects seem to contribute to hybrid incompatibilities not only in plants, but also in some animal systems (Wolf *et al*. 2014). As with the endosperm of flowering plants, genomic imprinting is an important feature of the mammalian placenta (Piedrahita 2011), and disrupted interactions between imprinted genes cause placental defects that result in abnormal hybrid growth in mice (Vrana *et al*. 2000). Although there has been much debate about the evolutionary mechanisms that account for the origin and maintenance of genomic imprinting in angiosperms and mammals (Haig and Westoby 1989, Kohler *et al.* 2012, Spencer and Clark 2014), most explanations involve some form of genomic coevolution. For example, the parental conflict theory posits that imprinting is shaped by conflict between males and females over maternal investment (Haig and Westoby 1989), and the coadaptation theory holds that integration of offspring and maternal genomes drives imprinted gene patterns (Wolf and Hager 2006). In plants, it has also been suggested that imprinted genes in the endosperm might evolve as an incidental by-product of a defensive silencing mechanism that targets invading foreign DNA in the embryo (Gehring *et al*. 2009, Hsieh *et al*. 2009). Whatever its evolutionary cause, genomic imprinting in the endosperm is highly variable among (Luo *et al.* 2011, Waters *et al.* 2011, Jiang and Kohler 2012) and even within (Waters *et al.* 2013, Pignatta *et al.* 2014) plant species. Thus, if divergence in patterns of imprinting does contribute to postzygotic reproductive isolation, it might be particularly important in the early stages of speciation.

Nevertheless, it is still unknown if parent of origin effects represent a major source of hybrid incompatibilities in flowering plants. Because F_1_ hybrid seeds combine divergent gene sequences, in addition to divergent patterns of imprinting, interactions among heterospecific alleles (*i.e*., Dobzhansky-Muller incompatibilities; Dobzhansky 1937, Muller 1942) at non-imprinted genes might also contribute to hybrid lethality. Indeed, in taxa without genomic imprinting (*e.g*., *Drosophila* and fish), there are many well-known cases of hybrid lethality due to incompatibilities between genes with bi-parental expression (*e.g*. Wittbrodt *et al*. 1989, Presgraves *et al*. 2003, Brideau *et al*. 2006, Tang *et al*. 2009). A key question, then, is whether F_1_ hybrid seed lethality in flowering plants is caused primarily by incompatibilities between imprinted or non-imprinted genes. The effects of the former are likely restricted to the endosperm, the site of most imprinting in plants, whereas the latter class of genes might act in the endosperm or in the embryo itself. Moreover, distinguishing between these two causes of F_1_ seed lethality is fundamentally important for understanding its evolution: imprinted genes represent a special class of loci that almost certainly reflect explicit genomic coevolution within species, rather than the independent fixation of alleles that only interact in hybrids.

Here we explore the mechanisms and genetics of reproductive isolation between two diploid species of *Mimulus*, *M. guttatus* and *M. tilingii*. In nature, these species are mostly allopatric, but they occasionally co-occur in high alpine areas. Because both species are primarily outcrossing with large, bee-pollinated flowers, sympatric populations might be expected to experience interspecific gene flow. However, early crossing studies reported that F_1_ hybrids between *M. guttatus* and *M. tilingii* are often difficult to generate (Vickery 1978), suggesting there might be some degree of postmating, prezygotic isolation that prevents fertilization and/or postzygotic isolation in the form of hybrid seed lethality. To investigate these possibilities, we began our study by intercrossing the two *Mimulus* species, characterizing both the mechanism and strength of reproductive isolation. In reciprocal crosses, we show there is only a modest reduction in hybrid seed number, but that almost all of these hybrid seeds are small, misshapen, and inviable.

This closely related *Mimulus* species pair thus presents a rare opportunity to test whether or not F_1_ hybrid seed lethality involves parent of origin effects. The finding that hybrid seed lethality is severe in *both* cross directions does not necessarily rule out reciprocal differences in its underlying genetic basis. That is, loci for *Mimulus* hybrid seed lethality might differ depending on their parent of origin. In this study, we use a powerful breeding design – backcrossing F_2_ hybrids reciprocally to each parent – to assess both the maternal and paternal contributions to *Mimulus* hybrid seed inviability. At the outset, our genetic analyses reveal two correlates of genomic divergence, transmission ratio distortion in F_2_ hybrids and chromosomal differentiation, which both provide insights into *Mimulus* speciation. Additionally, by performing high-resolution genetic mapping, we show that postzygotic reproductive isolation is primarily caused by loci with parent of origin effects, strongly suggesting a role for imprinted genes in the evolution of *Mimulus* hybrid seed lethality.

## MATERIALS AND METHODS

### Study system and plant material

The yellow monkeyflower *Mimulus guttatus* is highly polymorphic with natural populations distributed across much of western North America. The species occupies diverse environments, ranging from sand dunes along the Pacific coast to high alpine habitats. *M. tilingii*, a mat-forming perennial, occurs throughout much of the same geographic area, but is largely restricted to high elevations (2000+ m). Both species are self-compatible, but predominantly outcrossing with large, bee-pollinated flowers. *M. guttatus* and *M. tilingii* are closely related (Beardsley and Olmstead 2002) and belong to the same *Simiolus* section of primarily yellow-flowered taxa. Still, the two species have been classified as members of different species complexes (Vickery 1978) and variation at 16 nuclear loci suggest they form distinct genetic groups (Oneal *et al.* 2014). In areas of sympatry, there have been a few reports of putative hybridization (Lindsay and Vickery 1967, Carrie Wu pers comm.), but classic crossing experiments have shown that F_1_ hybrids are difficult to generate (Vickery 1978), suggesting reproductive isolation between *M. guttatus* and *M. tilingii* is strong.

In this study, we use one inbred line for each of the two focal species. The *M. guttatus* parental line (DUN10) is derived from a population located in the Oregon Dunes National Recreation Area along the Pacific coast. The *M. tilingii* parental line (LVR) originated from a high-alpine population in California’s Yosemite Valley (at 2751 m). Both of these inbred lines were formed by more than six generations of self-fertilization with single-seed descent.

### Measuring genomic divergence among *Mimulus* species

To determine the extent of genetic differentiation between *M. tilingii* and the well-studied *M. guttatus* complex (Brandvain *et al*. 2014), we measured genome-wide divergence (*i.e*., average pairwise nucleotide differences) among *M. tilingii*, *M. guttatus*, and *M. nasutus*. For *M. tilingii*, we generated new sequence data for the parental line LVR (see below) that yielded 68.4 million 150-bp paired-end reads. For *M. guttatus* and *M. nasutus*, we used six previously published lines (Table S1), including the parental *M. guttatus* line DUN10. Before alignment of *M. guttatus* and *M. nasutus* reads, we removed adapter contamination using Trimmomatic (Bolger *et al.* 2014) and confirmed removal using FastQC (www.bioinformatics.babraham.ac.uk/projects/fastqc/). We checked LVR for adapter contamination using FastQC but did not detect any. We aligned the paired-end reads from each line to the *M. guttatus* v2.0 reference genome (Hellsten *et al.* 2013) using the Burrows-Wheeler Aligner (BWA, BWA-MEM used for LVR, BWA-backtrack used for all others; Li and Durbin 2009) with a minimum quality threshold of Q29 (filtering done using SAMtools, Li *et al.* 2009). Following alignment, we removed PCR duplicates using Picard (http://picard.sourceforge.net/) and performed local realignment around indels using the Genome Analysis Toolkit (GATK, Depristo *et al.* 2011, McKenna *et al.* 2010).

We produced a high quality set of invariant sites and SNPs simultaneously using the GATK Unified Genotyper (Depristo *et al.* 2011, McKenna *et al.* 2010) with a site quality threshold of Q40. We excluded sites with more than two alleles and set cutoffs for coverage depth to a minimum of 10 and a maximum of 75 reads per site. To assign genotypes at heterozygous sites, we randomly selected one of two alternate alleles. We then quantified patterns of sequence variation for each pair of *Mimulus* samples using a custom python script (L. Flagel, pers. comm.). For each pairwise comparison, we counted the number of pairwise sequence differences and number of sites for which both samples have data above our quality and depth thresholds.

### Measurement of reproductive isolation and genetic crosses

To characterize postmating reproductive isolation between *M. guttatus* and *M. tilingii*, we performed crosses within and between the two parental lines (*N* = 20 each for DUN10 × DUN10, LVR × LVR, DUN × LVR, and LVR × DUN). For each of these crosses, we dissected one ripened fruit and measured total seed set. For each fruit, we also assessed seed viability by determining the proportion of viable seeds (number of viable seeds/total seeds). Seed viability was straightforward to score by eye: viable seeds were plump and tan in color, whereas inviable seeds were shriveled, darker in color, and often adhering to each other (Figure S1). To determine if our measure of seed viability is correlated with germination rate, we planted the seeds from a single fruit for crosses within parental lines (*N* = 2 each) and for a subset of our F_2_ × parental crosses (*N* = 20, see below). Because we found that our visual assessment of seed viability correlates strongly with germination rate (Spearman’s Correlation, *rho* = 0.92, *P* < 0.0001), all further measurements of seed viability were performed by eye.

To study the genetics of reproductive isolation between *M. guttatus* and *M. tilingii*, we intercrossed DUN10 (maternal parent) and LVR (paternal parent) to form F_1_ hybrids. We then backcrossed F_1_ hybrids (*N* = 20) reciprocally to each parental line and assessed seed viability as described above. To generate a recombinant population for genetic mapping, we self-fertilized a single F_1_ to form an F_2_ generation (*N* = 240). For each of these F_2_ hybrids, we performed reciprocal backcrosses to each parental line; seed viability was measured from a single fruit for each of the four cross treatments (Figure 1). Note that because the DUN10 line was used as the original maternal parent, all F_1_ and F_2_ hybrids carried an *M. guttatus* cytoplasm.

**Figure 1.**
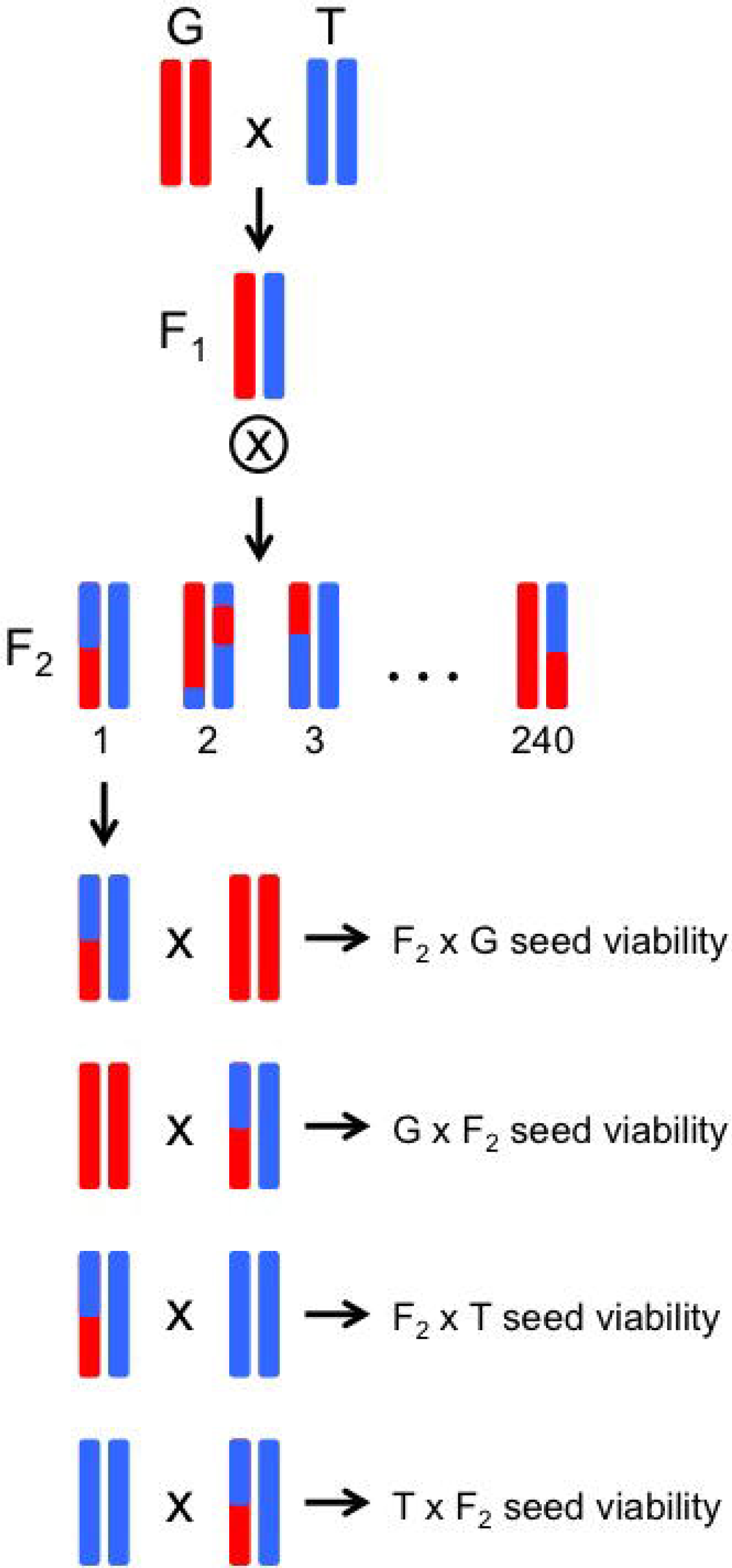
Crossing design to determine the genetic basis of seed lethality between *M. guttatus* and *M. tilingii.* We formed F_1_ hybrids by intercrossing *M. guttatus* (G, red) and *M. tilingii* (T, blue), and self-fertilized a single F_1_ to generate the F_2_ generation. We then reciprocally crossed F_2_ hybrids to each parental line and assessed seed viability. The diagram shows a single pair of chromosomes for each individualwith the maternal parent listedfirst.

All plants were grown using similar conditions at the University of Georgia. Seeds were planted into 2.5” pots containing Fafard 3b potting mix, chilled for 7 days at 4°C to promote germination, and then placed in a Conviron growth chamber with lights set to 16-hr days. Plants were bottom watered daily and temperatures were maintained at 22°C during the day and 16°C at night.

### DNA extraction, library preparation, and sequencing

For each of the 240 F_2_ individuals, we collected bud tissue into 96-well plates, immediately placed samples on dry ice, and stored them at -80°C. We isolated genomic DNA using a standard CTAB/chloroform extraction protocol as described in Holeski *et al.* (2014). Following extraction, we quantified DNA using the Quant-iT PicoGreen dsDNA Assay Kit (Invitrogen) and diluted each sample to 5 ng/*μ*L.

We generated a genomic library for genotyping the 240 F_2_ hybrids by using the multiplexed shotgun genotyping (MSG) method of Andolfatto *et al.* (2011). In 96-well plates, we digested 50 ng of genomic DNA (15 U *Mse*I; New England Biolabs) for each sample in 20-*μ*L reactions for 3 hr at 37°C followed by heat inactivation at 65°C for 20 min. Next, using 48 unique bar-coded adapters, we formed five sets of F_2_ individuals (each set with 48 F_2_s carrying bar-codes 1-48). We added 5 pmol of the bar-coded adapters to each well and performed ligation reactions using 1200 U of T4 DNA ligase (New England Biolabs) in total volumes of 50 *μ*L at 16°C for 3 hr followed by heat inactivation at 65°C for 10 min. We precipitated DNA in each well by adding 5 *μ*L of 3M sodium acetate and 50 *μ*L of isopropanol. We then pooled each set of 48 F_2_ individuals into a single tube, added 1 *μ*L of glycogen, and incubated overnight at 4°C. After resuspending each of the five samples in 100 *μ*L TE, we purified them using a 1.5 volume of Agencourt AMPure beads. Following purification, the Georgia Genomics Facility size-selected fragments of 375-425 bp using a QIAquick Gel Extraction Kit (QIAGEN). For each of the five sets of 48 F_2_ hybrids, we then performed 18 cycles of PCR using a Phusion High-Fidelity PCR master mix (New England Biolabs) and one of five unique “index” primers that bind to common regions in the bar-coded adapters. Following this PCR amplification, each of the 240 F_2_ individuals carried a unique bar-code/index identifier. We performed cleanup on each of the five sets using a 0.8 volume of Agencourt AMPure beads, resuspended in 40 *μ*L QIAGEN EB buffer, and quantified the DNA using a Qubit fluorometer. We combined the five indexed sets into one library of 240 F_2_ individuals at a final concentration of 15 *μ*M.

We sent our library of 240 F_2_ individuals to the Genome Sequencing Facility at Duke University for sequencing. We also included a sample of genomic DNA from the LVR *M. tilingii* line (the library was prepared by the Duke Genome Sequencing Facility). Because LVR is a highly inbred line, this genomic DNA should be nearly identical to that of the maternal line used to generate the F_2_ mapping population. Sequencing was performed across two lanes on an Illumina Hiseq 2500 for paired-end, 150-bp reads.

### Determining F_2_ genotypes

We used the MSG software pipeline (Andolfatto *et al.* 2011) to genotype 240 F_2_ hybrids between *M. guttatus* (DUN10) and *M. tilingii* (LVR). The MSG HMM algorithm requires reference genomes for both parents of the mapping population. To generate pseudo-reference sequences for DUN10 and LVR, we used the processed alignment files described above. We used SAMtools mpileup to call bases with a minimum base quality threshold of Q15 and depth capped at 250. We then used SAMtools bcftools followed by SAMtools vcfutils vcf2fq to output a fastq file for each line (with an indel filtering window of 20 bases, and minimum and maximum base depth thresholds set to five and the mean plus two times the standard deviation, respectively, for each line). Finally, we used Heng Li’s seqtk toolset (https://github.com/lh3/seqtk) to convert each fastq file to a fasta file, with a base quality threshold of Q15.

For all 240 *M.guttatus-M. tilingii* F_2_ hybrids, we assigned ancestry probabilities across chromosomes using the MSG v0.4.3 pipeline (Andolfatto *et al.* 2011). Because MSG does not consider reverse reads, we only used forward reads from the F_2_ hybrids. We removed low quality reads (less than an average Q6 across any 100-bp window) and reads with potential adapter contamination using Trimmomatic (Bolger *et al.* 2014). We then parsed reads for all 240 F_2_s by unique 6-bp barcode (within each of the five indices) using Stacks (Catchen *et al.* 2011). The parsed reads were aligned to each of the two parental reference genomes using bwa with default settings, followed by base calling with SAMtools. Next, a Hidden Markov Model (HMM) estimated a posterior probability of each possible ancestry (*i.e*., homozygous *M. guttatus*, heterozygous, or homozygous *M. tilingii*) across the genomes of each F_2_ individual. Finally, genotype probabilities were imputed for all F_2_ individuals at each informative genomic position that was typed in at least one individual. This process resulted in 151,669 SNPs across the 14 assembled chromosomes (SNPs per chromosome: mean = 10,834; min = 5,886; max = 19,217). The MSG parameters were set as follows: the priors on ancestry homozygosity for *M. guttatus* = 0.25, heterozygosity = 0.5, and homozygosity for *M. tilingii* = 0.25; the ancestry error probabilities for *M. guttatus* = 0.03 and for *A. tilingii* = 0.03; the genome-wide recombination rate = 28 crossovers per genome per generation (1 per chromosome for each F_1_ parent); and the arbitrary recombination scaling factor, rfac = 0.01 (qualitatively similar results were achieved at lower and greater values of rfac).

To facilitate linkage map construction, we converted ancestry probabilities from the MSG HMM to “hard ancestry calls,” assigning each individual a confident genotype at each marker. We first culled ancestry calls with less than 95% certainty by converting those calls to missing data. We then converted the remaining high confidence calls to a probability of 1.0 (or 0.0) for each parent. To remove markers with redundant information and convert the probabilities to hard calls, we used the script pull_thin.py (available on github at https://github.com/dstern/pull_thin) to thin markers with 100% identical neighbors (including identical missing data). Then, we thinned this dataset further using R/qtl (Broman *et al.* 2003) to include markers with at least one unique genotype across all 240 individuals (keeping markers with the least amount of missing data). The final, thinned dataset containing 2960 markers was then used for linkage map construction and MQM mapping, which requires hard genotype calls.

### Linkage mapping, transmission ration distortion, and QTL mapping

We performed linkage mapping in JoinMap 4.0 (Van Ooijen 2011) using a LOD threshold of 10.0, the Haldane mapping function, and the default Maximum Likelihood settings. In some cases, marker positions disagreed with their predicted locations from the *M. guttatus* v2.0 reference genome (often in regions known to be misassembled). For each linkage group, we assessed map quality using the nearest neighbor stress parameter in JoinMap, rlod scores output from the MSG pipeline, and visual inspection of genotypes to minimize double recombinants. Note that many of the regions that conflict between our genetic map and the genome assembly were also found in Holeski *et al.* (2014). Among the 240 F_2_ hybrids, six individuals were discovered to be pure *M. guttatus* (all markers were homozygous for DUN10 alleles); these six individuals were removed from all subsequent analyses. We also used JoinMap to test markers for significant non-Mendelian genotype frequencies.

To identify putative inversions between *M. guttatus* and *M. tilingii*, we compared genetic distances between the markers in our linkage map to physical distances from the *M. guttatus* v2.0 assembly. Because this final set of markers included only those with one or more unique genotype (among the 240 F_2_ hybrids), adjacent markers spanning large physical regions might represent areas of suppressed recombination. Alternatively, large physical jumps between adjacent markers could reflect low initial marker density due to a local paucity of SNPs. To exclude this second possibility, we measured initial marker densities in 500-kb windows using the raw, un-thinned marker outputs from the MSG pipeline. Regions identified as carrying putative inversions had initial marker densities similar to that of the genome-wide average.

We performed QTL mapping using MapQTL ®6 (Van Ooijen 2009). As a first step, we performed interval mapping (IM) using a 1-cM step size. Significance of QTL detected by IM was evaluated at the 5% significance level by permutation tests (*N* = 1000 permutations) at both the chromosome level and genome-wide level. Markers surrounding the QTL (i.e., with LOD scores exceeding the empirical threshold) were selected as cofactors for restricted Multiple QTL Mapping (MQM; analogous to composite interval mapping). To narrow the set of cofactors, we used the automatic cofactor selection tool in MapQTL with a threshold of *P* = 0.005. Automatic cofactor and restricted MQM analyses were repeated until a stable set of significant cofactors remained (note that restricted MQM excludes linked cofactors). At the end of this process, selected cofactors were used for a final round of restricted MQM to estimate for each QTL the maximum LOD, the additive genotypic effect (a), and the proportion of the F_2_ variance explained. As before, QTL were considered significant if LOD peaks reached the genome-wide and/or chromosome significance threshold of 5% (*N* = 1000 permutations). Finally, we calculated 1.5-LOD intervals for QTL found through MQM.

## RESULTS

### Nucleotide divergence among *Mimulus* species

Several recent studies have characterized population genomic variation within and between closely related species of the *M. guttatus* complex (Brandvain *et al*. 2014, Puzey and Vallejo-Marin 2014, Twyford and Friedman 2015), but samples from *M. tilingii*, which classic crossing studies suggest is more distantly related (Vickery 1978), were not included in these analyses. As a first step toward understanding the extent of genetic differentiation between these species, we examined genome-wide nucleotide variation within and between *M. guttatus*, *M. nasutus*, and *M. tilingii* (Table S2). Consistent with classic taxonomic groupings (Vickery 1978) and previous phylogenic work (Beardsley and Olmstead 2002), genome-wide divergence between *M. guttatus* and *M. tilingii* (*d* = 6.94%, *SE* = 0.28%) is substantially higher than divergence between *M. guttatus* and *M. nasutus* (*d* = 4.38%, *SE* = 0.09%). The latter pair is largely interfertile with ongoing introgression (Brandvain *et al.* 2014), and species divergence is comparable to levels of diversity within *M. guttatus* (π = 4.02%, *SE* = 0.19%). Higher genomic divergence between *M. guttatus* and *M. tilingii* suggests an earlier split for this pair and, potentially, stronger interspecific isolating barriers.

### Pattern of hybrid seed production and viability between *M. guttatus* and *M. tilingii*

To examine the strength of postmating reproductive isolation between *M. guttatus* and *M. tilingii*, we compared seed set from reciprocal interspecific crosses to that from crosses within each parental line (Figure S2). The DUN10 line of *M. guttatus* is a large flowered ecotype and produces significantly more seeds per fruit (mean = 313, *SE* = 16.6, *N* = 20) than the smaller flowered LVR line of *M. tilingii* (mean = 188, *SE* = 16.6, *N* = 20). For both sets of interspecifc crosses, seed set was lower than parental levels, but this reduction was only significant with *M. tilingii* as the maternal parent (*M. guttatus × M. tilingii*: mean = 167, *SE* = 16.6, *N* = 20; *M. tilingii × M. guttatus*: mean = 113, *SE* = 16.2, *N* = 20). Backcross seed set with *M. tilingii* as the maternal parent was similarly low (*M. tilingii ×* F_1_: mean = 103, *SE* = 19.2, *N* = 15). In contrast, seed sets from the other backcross classes were similar to that of *M. tilingii* (F_1_ × *M. guttatus*: mean = 166, *SE* = 18.6, *N* = 16; *M. guttatus ×* F_1_: mean = 214, *SE* = 17.5, *N* = 18; F_1_ × *M. tilingii*: mean = 204, *SE* = 18.0, *N* = 16). Note that because all F_1_ hybrids were generated using the DUN10 line of *M. guttatus* as the maternal parent, differences in seed set between reciprocal backcrosses to *M. tilingii* might involve cytonuclear interactions. Overall, there is a modest reduction in interspecific seed set, suggesting that reproductive isolating barriers (*e.g*. pollen-pistil incompatibilities) might partially interfere with the production of F_1_ hybrid seeds.

A much more dramatic isolating barrier was observed when we compared the proportion of viable seeds produced within and between species (Figure 2). Although seed viability differed significantly among parental lines (*M. guttatus*: mean = 0.95, *SE* = 0.02, *N* =20; *M. tilingii*: mean = 0.56, *SE* = 0.02, *N* = 20), both had much higher proportions than either interspecific cross. Indeed, F_1_ hybrid seed viability – in both directions of the cross – was <2% of the mid-parent value and not significantly different from zero (*M. guttatus × M. tilingii*: mean = 0.004, *SE* = 0.02, *N* = 20; *M. tilingii × M. guttatus*: mean = 0.01, *SE* = 0.02, *N* = 20). This F_1_ seed lethality represents an extremely strong barrier to interspecific reproduction, allowing less than 1% of F_1_ hybrid seeds to survive.

**Figure 2.**
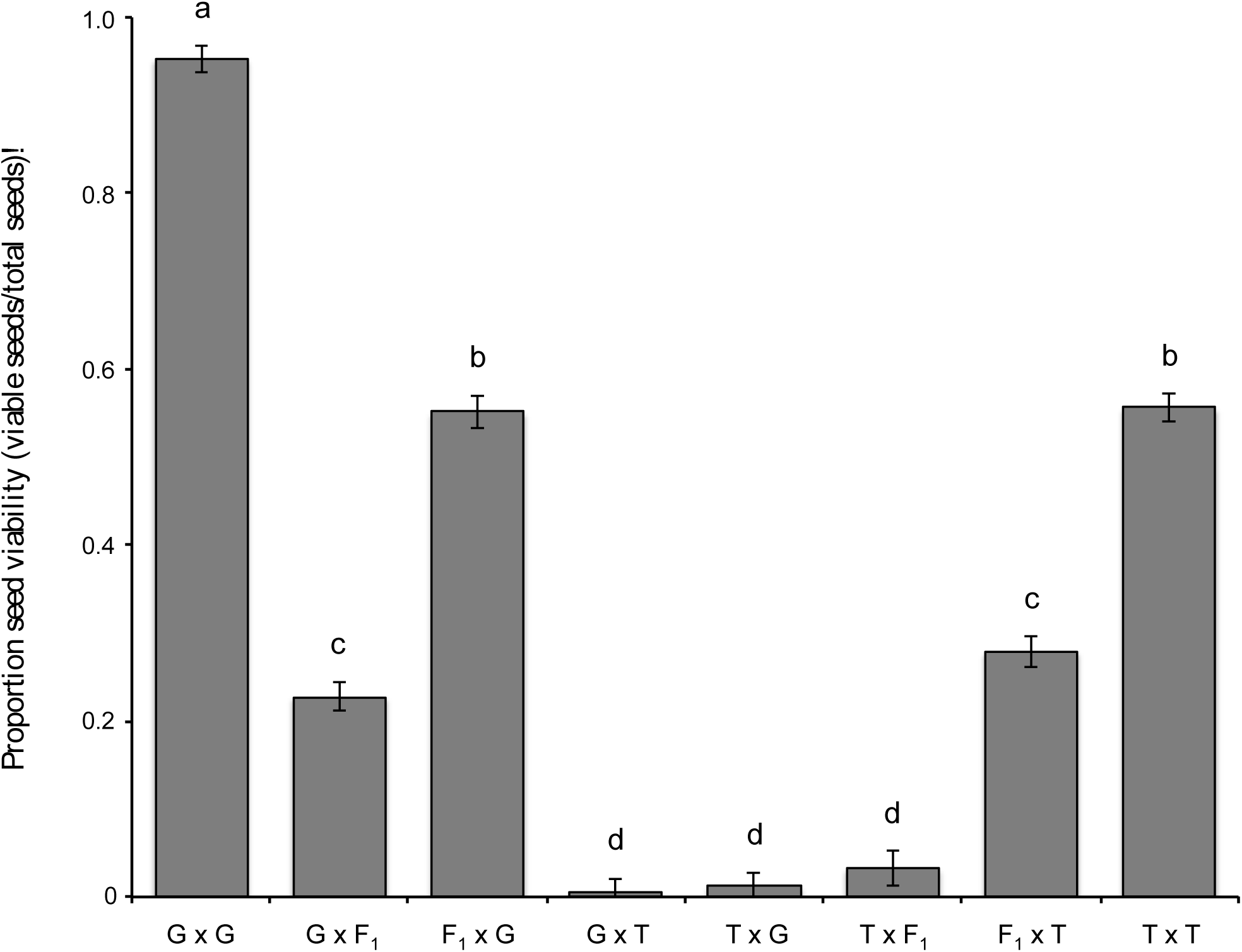
Mean seed viability varies among experimental crosses. Seed viability is highest from crosses within parental lines (*M. guttatus*: G × G, *M. tilingii*: T × T) and lowest in reciprocal interspecific crosses (G × T and T × G). Seed viability is generally intermediate in the four classes of backcross progeny (G × F_1_, F_1 ×_ G, T × F_1_, F_1 ×_ T). For all crosses, the maternal parent is listed first. Bars indicate standard errors and lower case letters above show significant differences by Tukey’s HSD test.

Moreover, this strong F_1_ hybrid lethality appears to be driven, at least in part, by loci with parent of origin effects: in backcrosses to either of the recurrent parents, hybrid lethality is much more severe when the F_1_ acts as the paternal parent. For the backcross to *M. guttatus*, this effect cannot be explained by cytonuclear interactions because the reciprocal backcross progeny inherit the same DUN10 cytoplasm. In contrast, the progeny of reciprocal backcrosses to *M. tilingii* do carry different cytoplasms; this fact, along with the possible maternal effects from *M. tilngii*, might explain the particularly low levels of seed viability observed when *M. tilingii* acts as the seed parent (in Figure 2, compare seed viability for T × F_1_ versus F_1_ × T). For the other backcross classes, seed viability is generally additive, with levels of seed failure among the progeny intermediate to that from parental and interspecific crosses (*e.g*., in Figure 2, compare seed viability for F_1_ × G to G × G and T × G).

### Transmission ratio distortion in the F_2_ hybrid mapping population

Our genetic map is based on 2960 markers and has a total length of 1318 cM with an average spacing of 0.4 cM. Two-thirds (1,987) of the markers genotyped in our F_2_ mapping population deviate from the expected 1:2:1 genotype ratios at α = 0.05, and nearly one-third (940) show significant transmission ratio distortion at a higher threshold (α = 0.001). The bias we observe is highly directional. Of the 940 markers distorted at α = 0.001, 692 have an excess of *M. guttatus* (GG) homozygous genotypes and a deficit of *M. tilingii* (TT) genotypes, whereas only 70 markers show the opposite pattern (excess of TT, deficit of GG). The vast majority of these markers (749 of 762) also show a significant bias in allele frequency from the expected 1:1 ratio (α = 0.05). The remaining markers with significant genotypic distortion (178) show an excess of heterozygotes, whereas no markers show a deficit of heterozygous genotypes.

Transmission ratio distortion is highly variable across the genome and affects 12 of the 14 linkage groups (Figure 3). By examining the genome-wide distribution of transmission bias, we identified 16 regions that contain genetically linked clusters of distorted markers. Ten of these regions show an excess of *M. guttatus* genotypes, three show an excess of *M. tilingii* genotypes, and three have an excess of heterozygotes (red, blue, and purple horizontal bars, respectively, in Figure 3). Two linkage groups (LG2 and LG6) have highly distorted markers across their entire lengths – both are overrepresented for *M. guttatus*.

**Figure 3.**
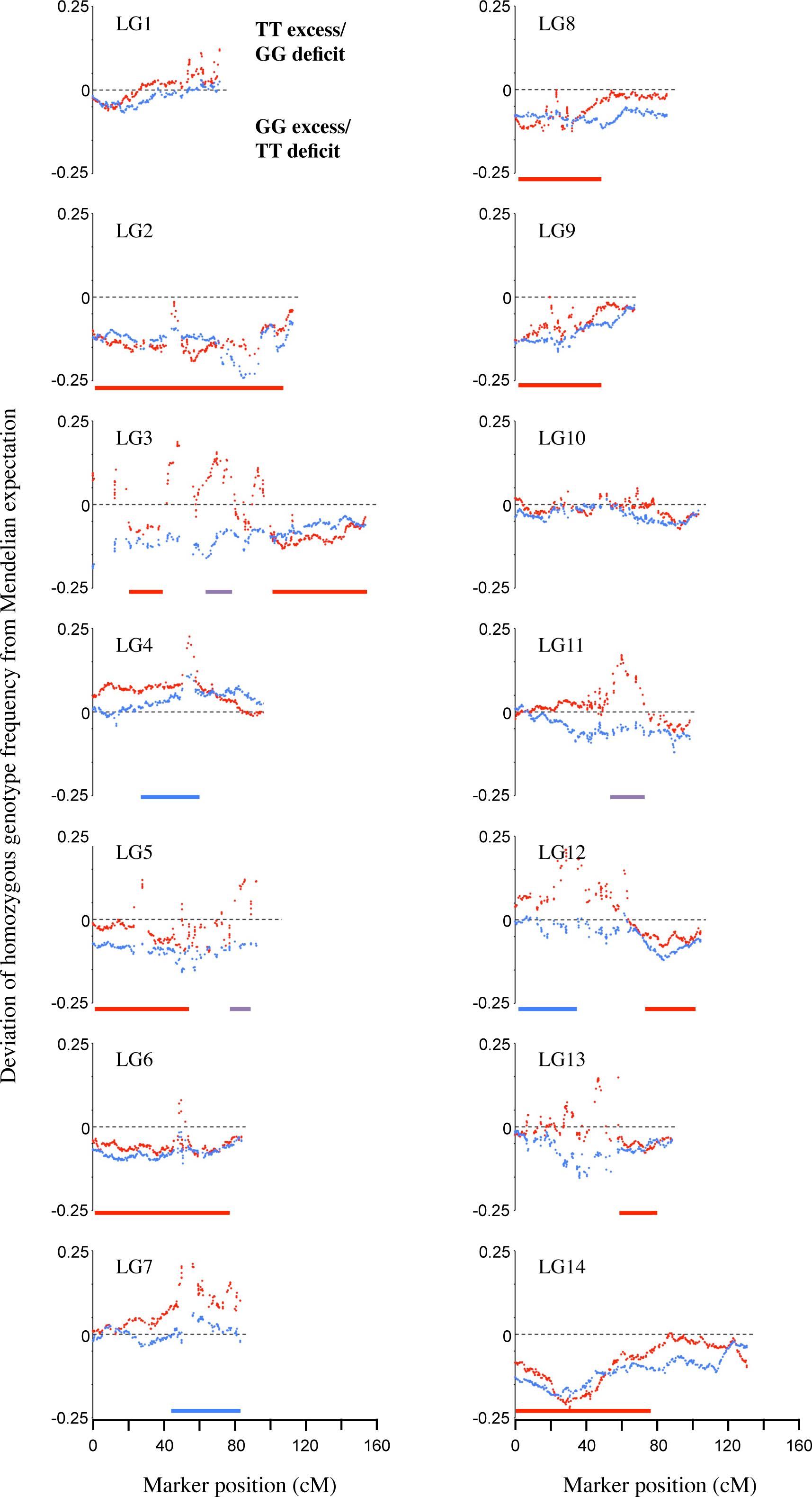
Transmission ratio distortion across the *M. guttatus × M. tilingii* linkage map. For each of the 14 linkage groups, homozygous parental genotypes at marker loci are shown in red (*M. guttatus*, GG) and blue (*M. tilingii*, TT). For each marker, the vertical positions of the red and blue dots show the deviations of genotype frequencies from the Mendelian expectation of 0.25. Note that the *M. tilingii* homozygote (TT) deviations are plotted directly [deviation = (frequency of TT) – 0.25] and the *M. guttatus* (GG) deviations are plotted as negative [deviation = – ((frequency of GG) – 0.25)]. As a result, the values above the zero line show excesses of TT homozygotes or deficits of GG homozygotes and values below the zero line show excesses of GG homozygotes or deficits of TT homozygotes. Horizontal bars beneath linkage groups indicate regions with excess GG (red), TT (blue), or heterozygotes (purple).

### Evidence for chromosomal inversions

Each of the 14 linkage groups in our genetic map corresponds to a chromosome from the *M. guttatus* v2.0 sequence assembly (http://www.phytozome.org), which represents ∼294 Mb of the genome (total haploid genome size is ∼450 Mb, http://www.mimulusevolution.org). Using genetic distances from our map and physical lengths from the genome sequence, the average recombination rate per chromosome is 4.6 cM/Mb (range = 3.4-8.1 cM/Mb, σ = 1.2 cM/Mb). The genome-wide estimate of (1318 cM/450 Mb) ∼2.9 cM/Mb is substantially lower because it includes portions of the genome not included in the sequence assembly. These results are broadly consistent with previous genetic maps from *M. guttatus* (*et al.* 2014, Holeski *et al.* 2014).

Across our genetic map, we observed three regions where a large number of physically dispersed markers map to the same location (Figure 4, regions highlighted in yellow), suggesting the presence of inversions between *M. guttatus* and *M. tilingii*. One of these regions extends from ∼0.9-7.6 Mb on chromosome 8 and overlaps with a previously discovered inversion known as *DIV1* that differentiates *M. guttatus* annual and perennial ecotypes (Lowry and Willis 2010, Oneal *et al.* 2014, Twyford and Friedman 2015). Because the DUN10 *M. guttatus* parent is known to carry the “perennial” *DIV1* arrangement (Lowry and Willis 2010), our results strongly suggest that *M. tilingii* is collinear with annual forms of *M. guttatus*. The two other putative inversions are from ∼13.4-18.8 Mb on chromosome 5 and ∼15.9-20.5 Mb on chromosome 13. This region on chromosome 5 overlaps with a putative inversion segregating within *M. guttatus* (Holeski *et al.* 2014).

**Figure 4.**
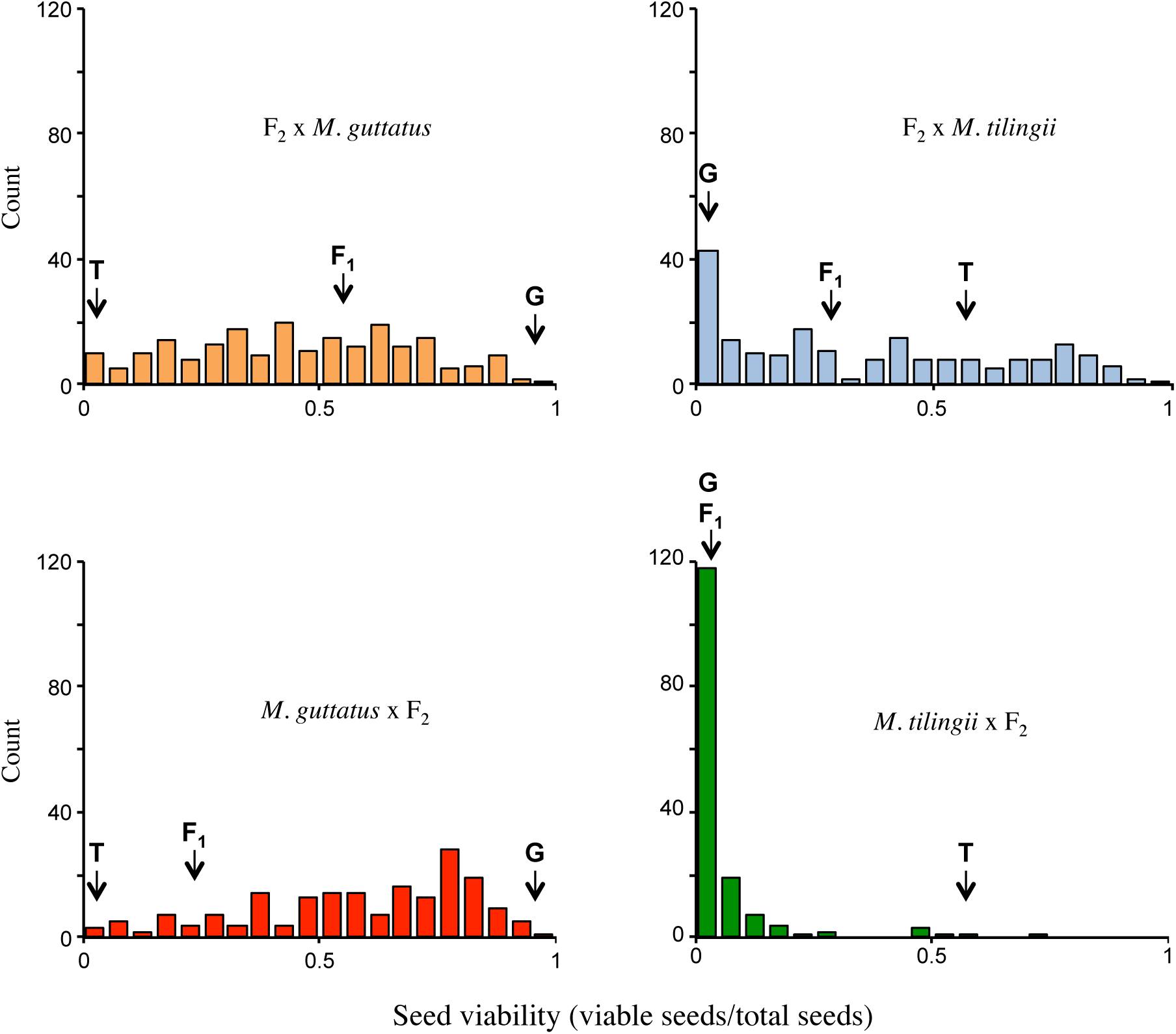
Frequency distributions of F_2_ hybrid seed viability phenotypes for the four experimental crosses. F_2 ×_ *M. guttatus* (average = 0.45, *N* = 215), *M. guttatus ×* F_2_ (average = 0.58, *N* = 190), F_2 ×_ *M. tilingii* (average = 0.36, *N* = 207), and *M. tilingii ×* F_2_ (average = 0.05, *N* = 158). Approximate values for parental (*M. guttatus*: G, *M. tilingii*: T) and F_1_ means are indicated with arrows in each histogram.

### QTL mapping of hybrid seed lethality and characterization of parent of origin effects

The distribution of hybrid seed viability varies substantially among the four different F_2_ cross types (Figure 4). Intriguingly, the distinct patterns of phenotypic variation in reciprocal crosses (F_2_ × G vs. G × F_2_; F_2_ × T vs. T × F_2_) suggest parent of origin effects on hybrid seed viability (but note that in cross types involving *M. tilingii* we cannot rule out an effect of cytoplasm). In three of the four cross types, seed viability appears generally consistent with additivity, with the means for F_1_ and F_2_ hybrids similar to the midparent values (see the first three panels of Figure 4). In contrast, seed viability from the *M. tilingii ×* F_2_ crosses is skewed strongly to the left, potentially suggesting a role for cytonuclear interactions and/or maternal effects. Additionally, we observed transgressive inheritance for seed viability in the F_2_ × *M. tilingii* crosses (with a substantial number of F_2_ hybrids producing more viable seeds than pure *M. tilingii*).

Our QTL analyses show a polygenic basis for hybrid seed lethality between *M. guttatus* and *M. tilingii*. For three of the four cross types, we detected multiple QTLs (18 total; Figure 5, Table 1). In the *M. tilingii ×* F_2_ cross, we found only a single QTL, but our power of detection was likely limited by a smaller sample size (only 158 F_2_ hybrids were phenotyped for this cross type) and non-normal distribution of seed viability (Figure 4; Beavis 1998). For 17 of the 18 QTLs, additive effects were in the expected directions based on parental values. Individual QTLs often had strong effects on seed lethality: reductions in backcross seed viability for the “lethal” versus alternative homozygote ranged from 20-61% (Table 1). On average, the seed viability of F_2_ hybrids carrying a lethal genotype at one of the QTLs was reduced by 39%.

**Table 1.** Summary of seed lethality QTL peak locations, statistical significance (LOD score), additive effect estimates (*a*), and viability effects.

| Cross | Linkage group | Position | LOD | $a^a$ | Reduction in viability<sup>b</sup> |
| --- | --- | --- | --- | --- | --- |
| $F_2 \times M. guttatus$ | 2 | 70.53 | 3.62 | 0.078 | 0.42 |
|  | 5 | 33.88 | 2.74 | 0.066 | 0.35 |
|  | 8 | 58.39 | 5.00** | 0.080 | 0.42 |
|  | 10 | 64.05 | 3.51 | 0.069 | 0.37 |
|  | 13 | 58.35 | 5.59** | 0.093 | 0.50 |
|  | 14 | 21.23 | 5.14** | 0.131 | 0.61 |
|  | 14 | 72.46 | 6.74** | 0.120 | 0.49 |
| $M. guttatus \times F_2$ | 2 | 57.95 | 3.06 | 0.063 | 0.20 |
|  | 4 | 2.36 | 2.95 | 0.067 | 0.21 |
|  | 6 | 13.41 | 2.63 | 0.082 | 0.25 |
|  | 7 | 66.99 | 3.14 | 0.086 | 0.26 |
|  | 13 | 51.32 | 3.31 | 0.089 | 0.27 |
| $F_2 \times M. tilingii$ | 2 | 72.73 | 2.93 | 0.092 | 0.41 |
|  | 6 | 31.94 | 3.91 | -0.094 | 0.41 |
|  | 7 | 20.17 | 3.09 | -0.073 | 0.33 |
|  | 9 | 7.73 | 3.17 | -0.156 | 0.45 |
|  | 14 | 78.10 | 7.57** | -0.140 | 0.55 |
| $M. tilingii \times F_2$ | 4 | 18.59 | 2.92 | -0.037 | 0.53 |

\*\*QTL is significant at a genome-wide threshold of 5% ( $N = 1000$ ). All other QTLs are significant at the chromosomal level with a threshold of 5% ( $N = 1000$ ).
<sup>a</sup>Positive values indicate the *M. guttatus* allele increases seed viability, negative values indicate the *M. tilingii* allele increases seed viability.
<sup>b</sup>Relative seed viability of the alternative homozygotes. The value reported is the reduction in backcross seed viability for the “incompatible” genotype relative to the “compatible” genotype.

**Figure 5.**
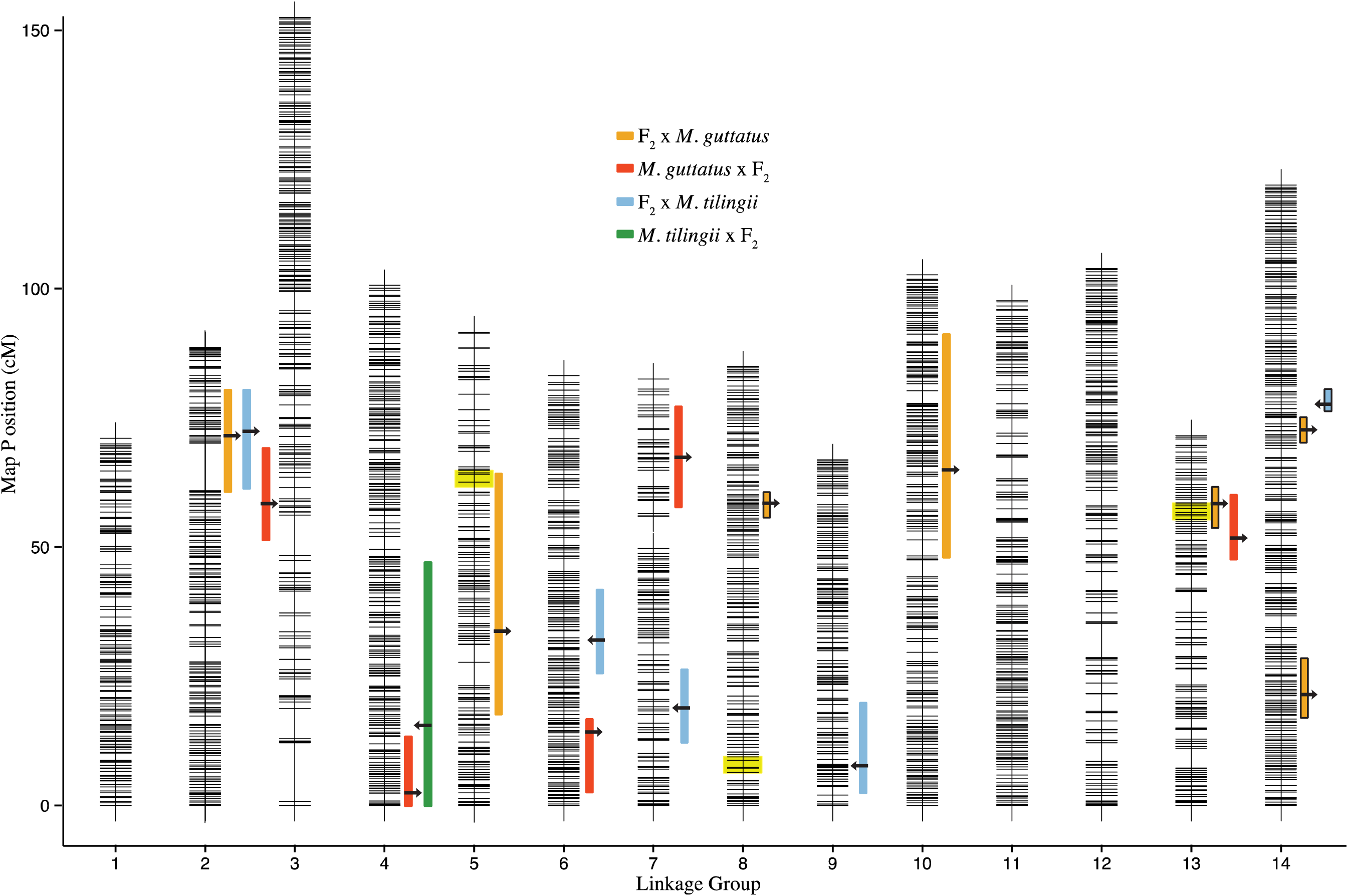
M. guttatus × M. tilingii genetic map and seed viability QTL. Each marker is shown as a horizontal line and the three hypothesized inversions are indicated by yellow shading. Arrows show the QTL LOD score peak locations and directions of additive effects (right pointing arrows indicate *M. guttatus* alleles increase the trait value, left pointing arrows indicate *M. tilingii* alleles increase the trait value). QTL bars show the 1.5 LOD drop confidence intervals and those outlined in black are significant at the genome-wide level (all others are significant at the chromosome level).

In crosses to both species, we discovered distinct sets of QTLs for hybrid seed lethality in reciprocal F_2_ crosses, implying that the allelic effects of these loci differ depending on whether they are inherited from the maternal or paternal parent. Indeed, there is little overlap between QTLs mapped in F_2_ × *M. guttatus* crosses and those mapped in *M. guttatus ×* F_2_ crosses (orange vs. red bars in Figure 5); the same is true for QTLs mapped in the two cross types involving *M. tilingii* (blue vs. green bars in Figure 5). Additionally, two QTLs show opposite allelic effects – in the predicted directions – when crossed to different species. One of these QTLs (on chromosome 4) affects hybrid seed lethality when it segregates in the paternal parent. The other QTL (on chromosome 14 at ∼72-78 cM) affects seed lethality through the maternal parent. Taken together, these results provide strong evidence that QTLs with parent of origin effects play a major role in *M. guttatus-M. tilingii* hybrid seed lethality.

One exception to this general pattern is the QTL on chromosome 2. At this locus, the *M. guttatus* allele increases seed fertility in three of the four cross types irrespective of direction or species. This result suggests that the chromosome 2 QTL is part of a hybrid incompatibility or simply reflects the segregation of deleterious alleles from the *M. tilingii* parent (in Figure 2, note the lower seed viability for this inbred line).

## DISCUSSION

In this study, we have shown that hybrid seed lethality is a highly effective isolating barrier between *M. guttatus* and *M. tilingii* with F_1_ seed viability being less than 1%. Additionally, we used a high-resolution mapping experiment to characterize the genetic basis of this widespread and exceptionally strong form of postzygotic reproductive isolation. Along with identifying hybrid lethality QTLs, our genetic analyses revealed both transmission ratio distortion (TRD) in F_2_ hybrids and chromosomal differentiation resulting in suppression of hybrid recombination. These phenomena are common in other plant mapping populations, and may provide insights into functional and genomic divergence between species. Furthermore, our reciprocal cross design allowed for direct tests of parent of origin effects on hybrid seed lethality at the QTL-level. Strikingly, most of the QTLs we identified contribute to hybrid seed lethality when inherited from only one of the parents (maternal or paternal), providing strong evidence for the involvement of imprinted genes. This finding is consistent with recent empirical work in *Arabidopsis* suggesting that imprinting plays a central role in triploid block (seed lethality in interploid hybrids), but our study is the first to identify both maternal- and paternal-effect loci for hybrid seed lethality between diploid species. Further work will be needed to identify the causal genes, but the results presented here suggest that divergence in genomic imprinting can generate strong postzygotic isolation between plant species in the early stages of divergence.

### Transmission ratio distortion and chromosomal differentiation between *Mimulus* species

In addition to QTLs for hybrid lethality, our mapping experiment revealed two common correlates of species divergence, transmission ratio distortion and evidence for chromosomal rearrangements. Just within the *M. guttatus* complex, TRD has been instrumental in the discovery of centromere-associated female meiotic drive (Fishman and Willis 2005, Fishman and Saunders 2008), loci underlying gamete competition and conspecific pollen precedence (Fishman *et al*. 2008), and potential cytoplasm-dependent hybrid incompatibilities (Lowry *et al*. 2009). In addition to these mechanisms, TRD in hybrids may also reflect inbreeding depression, barriers to fertilization, and postzygotic hybrid seed lethality. Similarly, suppression of recombination can reveal inversions or other rearrangements distinguishing species, which may be important in the development of both premating (Kirkpatrick and Barton 2006) and postzygotic barriers (Noor *et al*. 2001, Navarro and Barton 2003).

A large number of markers had distorted genotypic frequencies in our F_2_ mapping population, similar to other crosses within (Hall and Willis 2005) and between *Mimulus* species (Fishman *et al.* 2001, Fishman *et al.* 2015). The proportion of distorted markers in our F_2_ population (67% at α = 0.05, 32% at α = 0.001) is somewhat higher than the proportion found in an interspecific cross between *M. guttatus* and *M. nasutus* (49%, Fishman *et al.* 2001), consistent with the higher divergence between our focal species (*d* for *M. guttatus-M. nasutus* = 4.38%, *d* for *M. guttatus-M. tilingii* = 6.94%). In general, the distorted markers cluster in particular regions, suggesting that patterns of transmission distortion are caused by underlying loci, rather than by chance or error.

F_2_ mapping alone does not allow differentiation of the many mechanisms of TRD; however, our seedset data provide some clues about likely contributors. Least interesting, but a possibility in our cross, is inbreeding depression. Both parental lines were highly inbred, but they did not have equivalent fertility. Whereas the *M. guttatus* DUN10 line has high fitness, the *M. tilingii* LVR line produces a large fraction of inviable selfed seed (Figure 2). Segregation of deleterious recessive alleles fixed in the LVR parent could account, in part, to the genome-wide excess of *M. guttatus* genotypes and alleles we observe throughout the genome. Alternatively, TRD could reflect genetic interactions unique to hybrids. These heterospecific interactions can arise in gametes, where they have the potential to influence viability or fertilization success, or in F_2_ zygotes, where they might cause differential survival. Consistent with the action of gametic incompatibilities, in both reciprocal crosses of *M. guttatus* and *M. tilingii*, we observed a modest reduction in seed set (Figure S2), suggesting that certain loci might interfere with interspecific fertilization. Finally, the presence of strong F_1_ seed lethality, as well as the common observation of postzygotic hybrid incompatibilities between less divergent *Mimulus* species (*e.g*., Christie and MacNair 1987, Fishman and Willis 2006, Sweigart *et al*. 2007, Sweigart and Flagel 2015), suggests that Dobzhansky-Muller interactions causing F_2_ hybrid seed lethality might also contribute to TRD.

One intriguing possibility is that some of the same genetic loci might cause both hybrid seed lethality in our F_2_ crossing experiments and transmission ratio distortion in the F_2_ mapping population. At the LG2 QTL, F_2_ hybrids that carry *M. guttatus* alleles produce a higher proportion of viable seeds in three of four cross treatments. Similarly, if F_2_ seeds with *M. guttatus* alleles at LG2 are more likely to be viable themselves, it would lead to an overrepresentation of *M. guttatus* alleles in the F_2_ adults. The same effect might also occur at the distorted region on LG13; when the LG13 QTL carries *M. guttatus* alleles, it increases seed viability in reciprocal F_2_ crosses to *M. guttatus*. Interestingly, both the QTL and transmission ratio distortion on LG13 map to a putative inversion (Figure 5), suggesting that this region might contain multiple loci contributing to these phenotypes.

Importantly, for understanding the history of divergence between *M. guttatus* and *M. tilingii*, and for further studies of adaptation and speciation in this system, our genetic mapping revealed strong suppression of recombination in the *DIV1* region on chromosome 8 (Figure 5). The *DIV1* inversion defines widespread annual and perennial ecotypes of *M. guttatus* and contains QTLs for flowering time and growth-related traits that differentiate them (Hall *et al.* 2006, Lowry and Willis 2010, Twyford and Friedman 2015). Our finding that the LVR strain of *M. tilingii* is not collinear with the perennial DUN10 parent suggests that the perennial arrangement might be evolutionarily derived, consistent with an observed reduction in genetic diversity in the *DIV1* region in *M. guttatus* perennials (Oneal *et al.* 2014, Twyford and Friedman 2015).

### Loci with parent of origin effects cause strong reproductive isolation between *Mimulus* species

At first glance, the finding that severe F_1_ hybrid seed lethality occurs in both reciprocal crosses between *M. guttatus* and *M. tilingii* seems contrary to the idea that imprinted genes are involved. Unlike many interploidy and interspecies crosses, which often show pronounced reciprocal differences in seed lethality (see Thompson 1930, Haig and Westoby 1991), we detected no parent of origin effects in these first-generation *Mimulus* hybrid seeds. So, if imprinted genes *do* cause F_1_ hybrid seed lethality, the lack of reciprocal differences points to an independent genetic basis for the phenotype in each cross direction, implying genetic changes have evolved in both *Mimulus* lineages. Indeed, this is exactly what our mapping of distinct maternally- and paternally-contributed QTLs reveals: many loci with parent of origin effects contribute to *Mimulus* hybrid seed lethality. Of course, a more detailed phenotypic characterization of *Mimulus* F_1_ hybrid seed lethality might also reveal reciprocal differences at the level of endosperm growth and/or development that are associated with these distinct sets of genetic loci.

We found a polygenic basis for hybrid seed lethality between *M. guttatus* and *M. tilingii*. Across the four F_2_ backcross treatments, we detected 18 QTLs, although several of these QTLs seem not to be independent. On chromosome 14, for example, two maternally contributed QTLs map to overlapping regions and have opposite allelic effects in backcrosses to *M. guttatus* and *M. tilingii*. The simplest explanation for this pattern is that one QTL causes both effects: F_2_ hybrids carrying *M. guttatus* alleles at the causal locus promote seed compatibility in crosses to *M. guttatus*, whereas *M. tilingii* alleles improve seed viability in crosses to *M. tilingii*. Similarly, on chromosome 4, overlapping paternal QTLs have opposite phenotypic effects in the two cross treatments (in the predicted directions), suggesting a common genetic basis. For two additional genomic regions – on chromosomes 2 and 13 – we detected overlapping QTLs from reciprocal crosses, implying that allelic effects at the underlying loci do not depend on parent of origin. For all remaining, non-overlapping QTLs, dominance relations and/or genetic background effects might make detection more likely in one backcross than in the other. Fortunately, our four-way crossing design maximizes the chances of mapping such QTLs despite these complicating effects. In total, we found eight QTLs with seed lethality effects only through the maternal parent, three QTLs with effects only through the paternal parent, and two QTLs with effects through both parents. Similar to these results in *Mimulus*, postzygotic barriers affecting hybrid seed survival between *Arabidopsis* species (Burkart-Waco *et al.* 2012) and between *Capsella* species (Rebernig *et al*. 2015) are also determined by many genetic loci. In *Arabidopsis*, epistasis among the causal loci indicates that hybrid seed incompatibility is controlled by a complex genetic network (Burkart-Waco *et al.* 2012). Thus, in three diverse systems, strong F_1_ postzygotic isolation seems to have evolved between closely related species due to the accumulation of multiple incompatibilities that combine to cause severe defects in hybrid seed development.

Given the high variability in patterns of genomic imprinting even among strains of *Arabidopsis* (Pignatta *et al.* 2014), information from other species may have limited predictive value for identifying candidate genes in our *Mimulus* seed lethality QTLs. However, as a first step toward identifying candidate genes for parent of origin effects in *Mimulus* hybrid seed lethality, we performed a blast search of the *M. guttatus* reference genome using genes previously identified as imprinted in the *A. thaliana* endosperm (Hsieh *et al.* 2009, Wolff *et al.* 2011, Pignatta *et al.* 2014). We blasted 438 maternally expressed and 150 paternally expressed *Arabidopsis* genes and recovered 303 and 92 best hits, respectively, from the *M. guttatus* genome (using an e-value cutoff of 1E-06). Among these 395 *M. guttatus* genes, 26 co-localize with maternal QTLs and six co-localize with paternal QTLs. Based on their annotations, none of these genes has an obvious functional role in endosperm development, and of course, there is no guarantee that any of them are imprinted in *Mimulus*. Thus, further fine-mapping and functional/expression analyses will be necessary to identify the molecular basis of parent of origin effects in Mimulus hybrids; such analyses will also enlarge our understanding of this highly dynamic phenomenon beyond a few model systems.

A key question is which evolutionary forces might have led to divergence at hybrid lethality loci between *M. guttatus* and *M. tilingii*. One intriguing possibility is that F_1_ hybrid lethality in both reciprocal crosses is the outcome of unique coevolutionary histories between imprinted genes and their targets in each of the two *Mimulus* lineages. According to the parental conflict model, maternally expressed genes should be selected to restrict endosperm growth, whereas paternally expressed genes might function to promote growth (Haig and Westoby 1989). If patterns of genomic imprinting have evolved in *Mimulus* because of parental conflict over maternal investment, it is certainly possible that different species have resolved this conflict using distinct genetic routes. In nature, both *M. guttatus* and *M. tilingii* are predominantly outcrossing, so parental conflict has the potential to be strong within each species (Brandvain and Haig 2005). Of course, lineage-specific changes may also occur if imprinted genes have evolved by some other selective mechanism [*e.g*., coadaptation between maternal and offspring traits (Wolf and Hager 2006)] or by incidental proximity to silenced TEs (Gehring *et al*. 2009, Hsieh *et al*. 2009). In the latter case, patterns of genomic imprinting might be expected to be particularly dynamic given the high degree of variation in TE position within and between plant species (Tenaillon *et al.* 2010, Cao *et al.* 2011). Interestingly, F_1_ seed lethality also occurs in some crosses between geographically distant populations of *M. tilingii* (Sweigart *et al.*, unpubl. results), suggesting that, as in *Arabidopsis* and maize (Waters *et al.* 2013, Pignatta *et al.* 2014), there may be variation within *M. tilingii* for patterns of genomic imprinting.

### Conclusions

In this study, we have presented the first investigation of reproductive isolation and its genetic basis between *M. guttatus* and *M. tilingii*. Both species have abundant natural populations that occasionally co-occur, potentially providing the opportunity for interspecific gene flow. However, we have shown that there is incredibly strong postzygotic reproductive isolation between these species. Our quantitative genetic analysis of hybrid seed lethality – the most comprehensive to date between diploid plant species – confirms a central role for genetic loci with parent of origin effects. These findings set the stage for future fine-mapping, phenotypic characterizations of hybrid seed development, and gene expression studies to identify the underlying genes. This possibility, along with the potential for discovering natural variation in genomic imprinting, makes this *M. guttatus-M. tilingii* system a particularly rich one for investigating the evolutionary mechanisms of this early acting form of postzygotic reproductive isolation.

## ACKNOWLEDGEMENTS

We thank Rachel Hughes for expert greenhouse care and lab assistance. Kelly Dyer, Rachel Kerwin, John Willis, and Matt Zuellig made thoughtful comments on an earlier draft of this paper, which greatly improved it. We are grateful to Adam Bewick for bioinformatics assistance, and to Molly Schumer and David Stern, who provided invaluable MSG programming advice. We also thank the Duke Genome Sequencing Core and the Joint Genome Institute for generating the sequences used in this study. This work was supported by a Center for Undergraduate Research Opportunities (CURO) Summer Fellowship at the University of Georgia to A.G.G. and a National Science Foundation (NSF) grant DEB-1350935 and to A.L.S.

